# Loss of the *PPE71-esxX-esxY-PPE38* locus drives adaptive transcriptional responses and hypervirulence of *Mycobacterium tuberculosis* Lineage 2

**DOI:** 10.1101/2025.04.16.649112

**Authors:** Benjamin Koleske, Courtney Schill, Saranathan Rajagopalan, Somnath Shee, Yazmin B. Martinez-Martinez, Manish Gupta, Jessica Shen, William R. Jacobs, William R. Bishai

## Abstract

*Mycobacterium tuberculosis* (*M.tb*) is remarkable for its immense global disease burden and low mutation rate. Despite strong selective pressure, *M.tb* shows frequent deletions at the *PPE71–38* locus, most notably in hypervirulent L2 Beijing strains. Here, we show that loss of the *PPE71– 38* locus causes increased stress response gene expression and increased triglyceride levels. In addition, we demonstrate that re-introduction of *PPE71* into the L2 strain HN878 suppresses the baseline elevation of these transcripts, while overexpression of *PPE71* increases the localization of PE_PGRS proteins and lipoproteins to the *M.tb* outer mycomembrane. Mouse infection confirmed the hypervirulence of the *PPE71–38* deletion strain and conversely showed that *PPE71* overexpression attenuates *M.tb*. Our results indicate that loss of *PPE71–38* is sufficient to drive an adaptive transcriptional response seen in *M.tb* L2 strains that likely contributes to the hypervirulence of this lineage.

## Introduction

Tuberculosis (TB) remains the single leading cause of infectious disease mortality, with approximately 10.8 million new cases and 1.25 million deaths reported globally in 2023 (1). The causative organism, *Mycobacterium tuberculosis* (*M.tb*), dates back to antiquity and possesses essentially no environmental or animal reservoir beyond its human host (2–4). Due to the overall slow mutation rate of *M.tb* and the absence of natural transformation or lateral gene transfer, *M.tb* exhibits greater clonality and reduced genetic diversity compared to many pathogenic bacteria (5–9). An ancient microbe with a single hostile niche would be expected to experience strong purifying selection, and indeed nonsynonymous mutations are underrepresented in many genes across *M.tb* isolates (2, 7–11). Conversely, the odd beneficial mutation often rapidly expands across clinical isolates, as is seen in drug resistance (12–15). However, many expanded genetic variants remain poorly understood even if their variations have been linked to increased pathogenesis (10, 16). Hence, there is a pressing need to examine the polymorphisms that naturally arise in *M.tb* isolates, particularly those that converge across many strains.

*M.tb* strains can be divided into 10 lineages (L1–L10) differentiated by large genomic regions of difference (17–21). Lineages 2–4 comprise the “modern” *M.tb* lineages and cause TB disease broadly across the globe, while the remaining lineages are limited to smaller, specific geographies (22, 23). Correspondingly, L2–4 are monophyletic and share a deletion, TbD1, that promotes bacterial survival during hypoxic and oxidative stress (3, 24, 25). Subsequently, additional acquired polymorphisms have rendered the L2 strains, also termed the Beijing strains, particularly hypervirulent. The L2 lineage has been associated with greater transmissibility, worse clinical outcomes, and increased development of drug resistance mutations (26–29).

Curiously, L2 strains have a high frequency of gene deletions and perturbations at a locus containing two nearly identical PPE genes (named for conserved Pro-Pro-Glu residues), *PPE71* and *PPE38* (30–32). While *PPE71–38* disruption is most frequent in L2, it is also seen sporadically in other *M.tb* lineages as well as several animal-adapted *M.tb* complex strains (30–32). The non-overlapping nature these genetic lesions indicates that they arose several times independently, although it is unclear why there is apparent pressure to lose these *PPE* genes.

This convergent loss-of-function is complicated by existing knowledge on the role of the *PPE71–38* locus, which is required for the secretion of PE_PGRS proteins (33, 34). Among mycobacterial type VII secretion system substrates, PE_PGRS proteins are a diverse, evolutionarily recent family that appends a highly repetitive polymorphic GC-rich repeat sequence (PGRS) to the PE (Pro-Glu) N-terminal secretion domain (35–37). Type VII secretion system substrates are highly upregulated by *M.tb* during host infection, with PPE71–38 and PE_PGRS proteins among the most abundant bacterial proteins detected in infected guinea pig lung tissue at 30 and 90 days post-infection (38). Thus, PE_PGRS proteins appear to benefit the pathogen, and indeed virulence-promoting functions have been proposed for several PE_PGRS family members (39–41). Paradoxically, loss of *PPE71–38* and thus PE_PGRS secretion does not impair mycobacterial pathogenicity but rather appears to exacerbate virulence (33, 42). Therefore, we sought to understand why *PPE71–38* locus deletion improves bacterial fitness.

To generate strains with a range of *PPE71–38* gene doses, we deleted the *PPE71–38* locus and constructed complement and overexpressor addback strains in the *M.tb* H37Rv background. We infected these strains into BALB/c mice and recapitulated the known hypervirulence of the *PPE71–38* deletion strain (33). To identify the molecular basis of this hypervirulence, we performed transcriptomic and proteomic assessments on our panel of strains. From this, we identified that *PPE71–38* deletion caused *M.tb* to upregulate stress response and carbon metabolism genes and to increase triglyceride levels. In contrast, increased expression of *PPE71–38* boosted the abundance of lipoproteins in the *M.tb* cell wall. Finally, we expressed *PPE71* in the L2 strain HN878 (43), which naturally lacks expression of the *PPE71–38* locus, and demonstrated that restored *PPE71* levels were sufficient to repress the transcriptional responses we observed with *PPE71–38* deletion. Taken together, our results suggest that the *PPE71–38* locus disruption seen in L2 *M.tb* strains may allow the organism to adapt pre-emptively to stress conditions in the host environment and may account for the hypervirulence phenotype of the Beijing lineage.

## Results

### *PPE71* locus deletion and addback in *M.tb* H37Rv

To investigate the consequences of *PPE71–38* locus deletion, we used a two-step specialized phage transduction method to make an unmarked deletion of *PPE71*, *esxX*, *esxY*, and *PPE38* in *M.tb* H37Rv and designated this strain Δ7138 (**Fig. 1A**). Next, we transformed the Δ7138 strain with *PPE71*, *PPE71-esxX-esxY*, and *PPE71-esxX-esxY-PPE38* complementation plasmids to generate the Δ7138+71, Δ7138+71XY, and Δ7138+71XY38 strains, respectively. In addition, we transformed both WT H37Rv and Δ7138 with the empty pMH94 vector and with the PPE71 overexpressor construct to generate WT+vector, WT+71OE, Δ7138+vector, and Δ7138+71OE strains.

**Figure 1:**
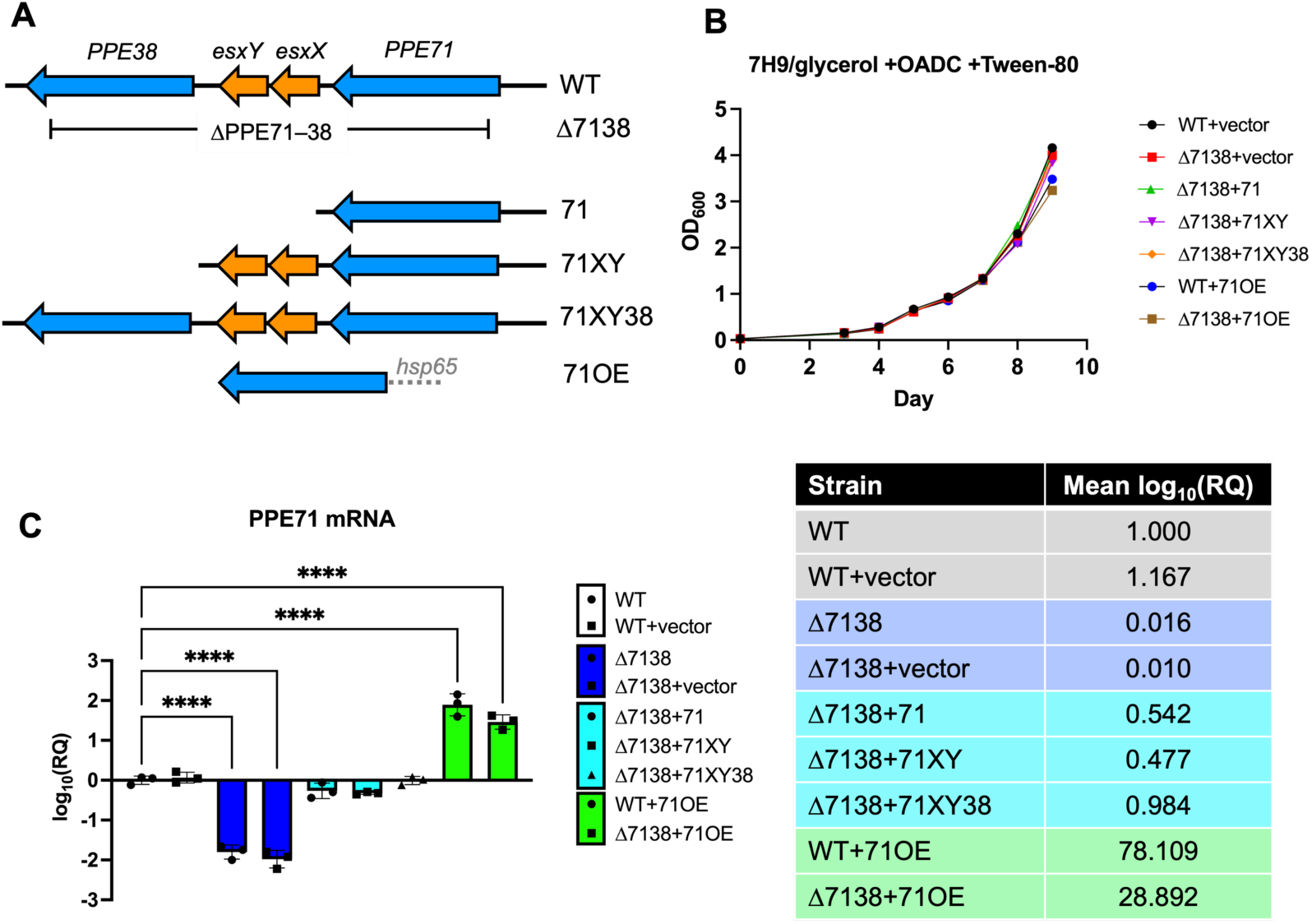
Design and validation of H37Rv Δ7138 deletion strain and addbacks. (A) Schematic diagram of the wild-type *M.tb* H37Rv *PPE71–PPE38* locus and the location of the Δ7138 deletion. Constructs to add back portions of the locus are shown below. All addback constructs use the native *PPE71* promoter aside from 71OE, which uses the *hsp65* promoter from *M. leprae* to achieve constitutive overexpression. (B) Growth curves of *M.tb* WT, deletion, and addback strains in 7H9 broth supplemented with 0.5% glycerol, 10% OADC, and 0.05% Tween 80. (OD_600_: optical density at 600 nm.) (C) Relative expression (log_10_ fold-change) by qPCR of *PPE71* mRNA in *PPE71* deletion and addback strains normalized to the parental WT H37Rv strain. *M.tb* 16S rRNA was used as an internal control. Mean relative expression values are provided in the accompanying table. (mean±SD; n=3; ****: p<0.0001 by one-way ANOVA.)

To assess the necessity of the *PPE71*–*38* locus for *in vitro* growth, we compared the growth kinetics of our panel of *PPE71* deletion and addback strains. All strains displayed little to no difference from the WT+vector strain when grown in complete 7H9 medium for 9 days, measured by optical density at 600 nm (OD_600_) (**Fig. 1B**). We assessed the expression of *PPE71* mRNA in these strains and found a >50-fold reduction from WT in the Δ7138 strain background and a >25-fold elevation from WT in strains bearing the 71OE construct (**Fig. 1C**). Hence, *in vitro* growth of these strains in complete medium was minimally affected despite pronounced changes in *PPE71* transcript levels.

### Loss of the *PPE71–38* locus confers hypervirulence with increased lung immunopathology

Since *PPE71* gene dose had a minimal effect on *M.tb* growth *in vitro*, we next infected our panel of *PPE71* strains into BALB/c mice model by aerosol infection (**Fig. 2A**). Lung inocula assessed by Day 1 lung colony-forming units (CFUs) demonstrated no significant differences between initial implants (**Fig. S1A**). We observed an increase in lung CFUs at Week 4 in the Δ7138+vector strain compared to WT+vector or WT+71OE (**Fig. 2B**). However, these differences were no longer present in the lungs at Week 8, although there was a trend towards the Δ7138+vector strain having the highest CFU counts (**Fig. 2D**). We also measured spleen CFUs during these same timepoints and found no significant differences between these strains (**Fig. S1B–C**).

**Figure 2:**
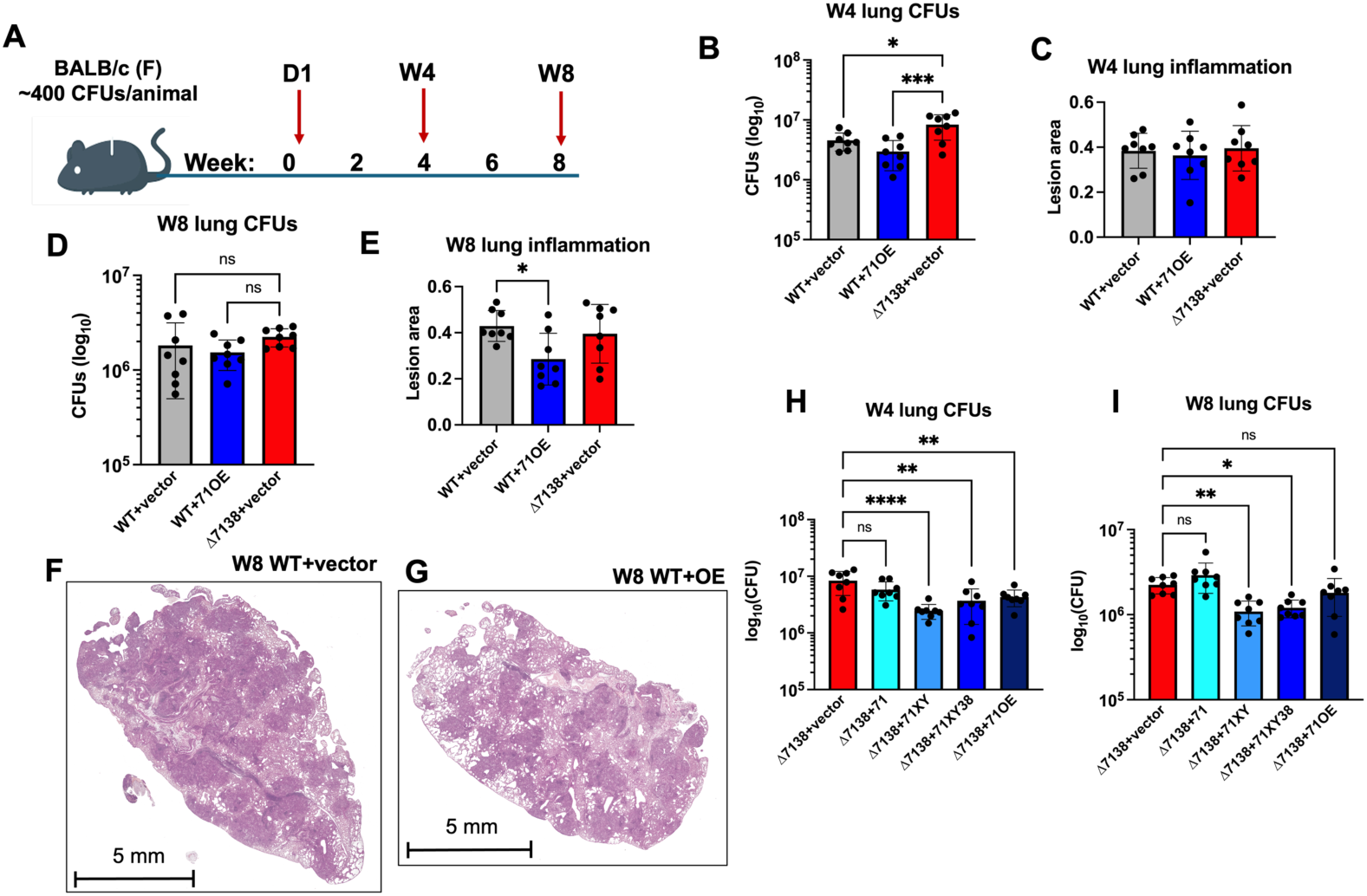
H37Rv Δ7138+vector is hypervirulent in a mouse model, while WT+71OE is attenuated. (A) Design for aerosol infection of *M.tb* H37Rv *PPE71* variant strains into female BALB/c mice. Animals were sacrificed on Day 1 to confirm inocula and on Week 4 and Week 8 as experimental timepoints. (B) Lung CFUs from the Week 4 timepoint for WT+vector, WT+71OE, and Δ7138+vector groups. Comparisons by one-way ANOVA compared to Δ7138+vector are depicted. (n=8 each, *: p<0.05; ***: p<0.001.) (C) Proportion of the lung occupied by lesions at Week 4 from WT+vector, WT+71OE, and Δ7138+vector groups, as quantified from hematoxylin & eosin (H&E) images. No significant differences were noted by one-way ANOVA. (n=8 each.) (D) Lung CFUs from the Week 8 timepoint for WT+vector, WT+71OE, and Δ7138+vector groups. Comparisons by one-way ANOVA compared to Δ7138+vector are depicted. (n=8 each, ns: non-significant.) (E) Proportion of the lung occupied by lesions at Week 4 from WT+vector, WT+71OE, and Δ7138+vector groups, as quantified from H&E images. Significant comparisons by one-way ANOVA are depicted. (n=8 each, *: p<0.05.) (F–G) Representative H&E histology images from the Week 8 WT+vector (F) and Week 8 WT+OE (G) groups, demonstrating reduced lesion area in the lung infected with WT+OE at this timepoint. (H–I) Lung CFUs from Week 4 (H) and Week 8 (I) timepoints for groups with *PPE71* addback strains in the Δ7138 background. Comparisons by one-way ANOVA compared to Δ7138+vector are depicted. (n=8 each, ns: non-significant; *: p<0.05; **: p<0.01; ****: p<0.0001.)

Beyond CFU counts, the immunopathology elicited by different *M.tb* strains is an important virulence parameter. Consequently, we quantitatively evaluated H&E images of infected mouse lung tissue at Week 4 and Week 8 for extent of inflammation. We did not find any significant differences in percent lung lesion area between WT+vector, WT+71OE, or Δ7138+vector strains at Week 4 (**Fig. 2C**). However, we found that the WT+71OE strain produced less inflammatory infiltrate at Week 8 than WT+vector (**Fig. 2E–G, Fig. S2**). This finding suggests attenuation of the WT+71OE strain at the later timepoint.

Next, we evaluated the effect of progressive *PPE71* gene dose addback into the Δ7138+vector strain. At Weeks 4 and 8, we found that adding *PPE71* under its native promoter alone (Δ7138+71) was insufficient to suppress hypervirulence (**Fig. 2H–I**). However, restoring *esxX* and *esxY* alongside *PPE71* (Δ7138+71XY) or the full *PP71–38* locus (Δ7138+71XY38) was sufficient to significantly decrease bacterial lung CFU burdens. Addition of overexpressed *PPE71* (Δ7138+71OE) likewise decreased virulence at Week 4, though this effect was lost by Week 8.

### PPE71 localizes to the outer mycomembrane of *M.tb*

Previous work has established that the *PPE71–38* locus is required for export of PE_PGRS family proteins from *M.tb* (33). PE_PGRS family proteins are known to be exported through the inner mycomembrane by the ESX-5 secretion system as a first step in secretion; however, it remains unknown how they traverse the waxy outer layer of the mycomembrane (34, 44). We reasoned that, if PPE71 is involved in this secondary step of PE_PGRS protein transport, then it may localize to the outer mycomembrane.

To detect PPE71 by Western blotting, we used a novel antibody generated against a predicted C-terminal epitope of PPE71. While this polyclonal antibody showed non-specific immunoreactivity with a ∼50 kilodaltons (kDa) protein present in mycobacterial lysates, it demonstrated a specific band in whole-cell lysate from the 71OE-bearing strains near the predicted molecular weight of PPE71 (37 kDa) (**Fig. 3A**). To assess localization of PPE71, we incubated WT+vector and WT+71OE with proteinase K for 5-minute intervals from 0 to 15 minutes. Because the bacterial cell wall remained intact, intracellular proteins such as FtsZ were unaffected by the protease (**Fig. 3B**, lower panel). However, the PPE71 band in the WT+71OE strain was promptly lost after just 5 minutes of protease treatment, demonstrating that the C-terminal epitope of PPE71 is fully exposed to the extracellular environment (**Fig. 3B**, upper panel).

**Figure 3:**
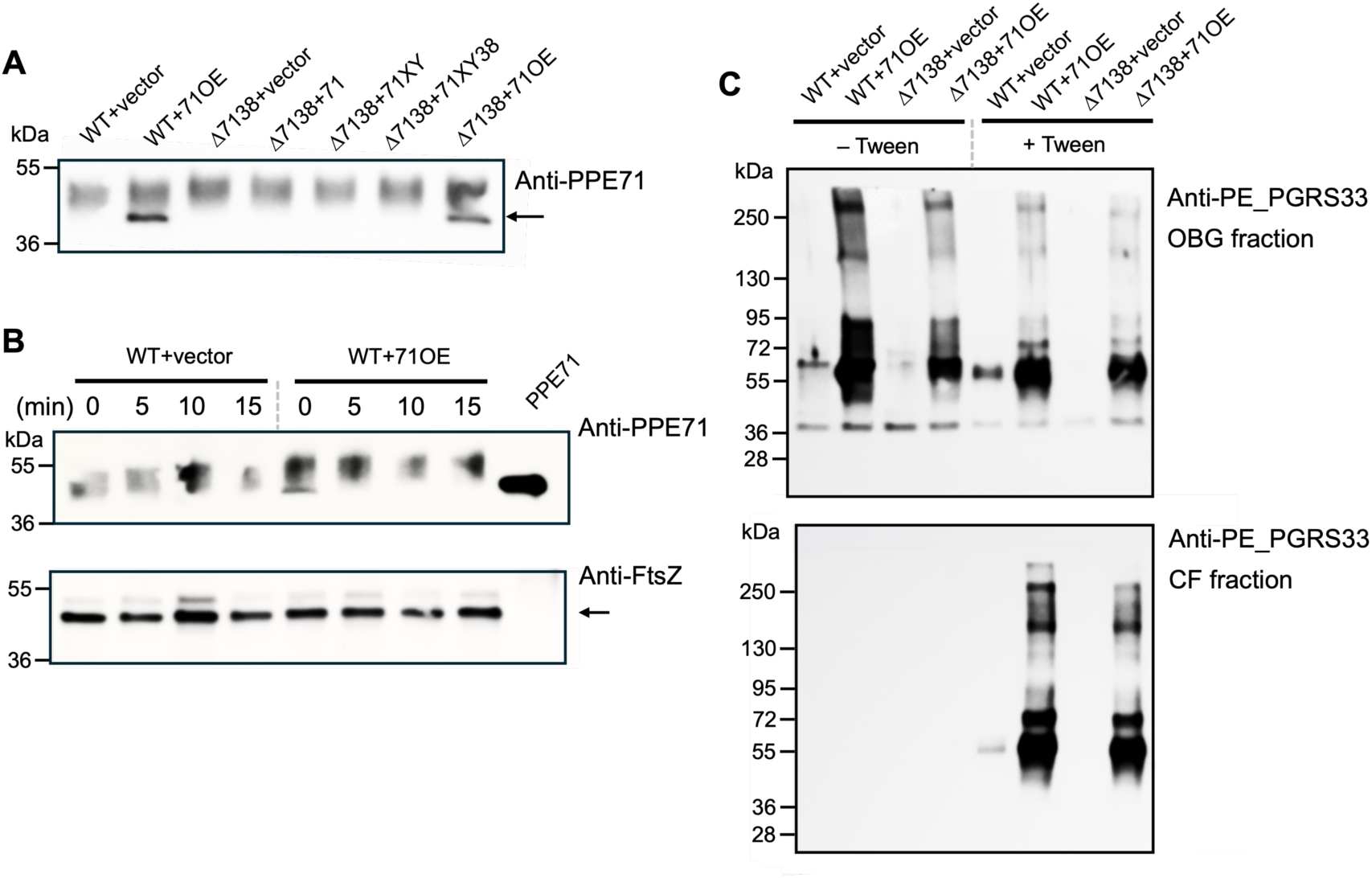
PPE71 is exposed on the *M.tb* cell surface and is needed to localize PE_PGRS proteins into the outer mycomembrane. (A) Western blot of whole-cell lysate from *M.tb PPE71* variant strains blotted with anti-PPE71 antibody. PPE71 protein is visible in the 71OE strains and is indicated with an arrow. (kDa: kilodaltons.) (B) WT+vector and WT+71OE cultures were incubated with proteinase K for the indicated time, then reactions were quenched. Whole-cell lysate was subjected to Western blotting with anti-PPE71 CTD (top) and anti-FtsZ (bottom; indicated with arrow). Recombinant purified PPE71 protein was included as a control (right lane). (kDa: kilodaltons.) (C) WT+vector, WT+71OE, Δ7138+vector, and Δ7138+71OE cultures were incubated overnight in complete 7H9 broth with or without Tween 80. Cell wall fractions obtained by octyl glucoside (OBG) incubation (top) and culture filtrate (CF) fractions (bottom) were subjected to Western blotting with anti-PE_PGRS33 antibody. (kDa: kilodaltons.)

### *PPE71* overexpression dramatically increases PE_PGRS protein secretion into the outer mycomembrane

Next, we aimed to confirm the relationship between *PPE71* gene dose and PE_PGRS secretion. In particular, we wondered whether overexpressing *PPE71* could enhance PE_PGRS secretion in excess of WT levels. To examine this, we used an antibody that targets the PGRS domain of PE_PGRS33 (44). Notably, this antibody has been shown to bind numerous other members of the PE_PGRS family, likely due to high sequence similarity (33, 34). To extract the outer mycomembrane, we incubated intact *M.tb* with n-octyl-β-D-glucoside (OBG), a nonionic detergent that achieves non-lytic solubilization of this mycobacterial outer membrane (45, 46).

Compared to the WT+vector strain, we found that both WT+71OE and Δ7138+71OE hypersecreted multiple PE_PGRS proteins (**Fig. 3C**). Minimal PE_PGRS proteins were detected in the Δ7138+vector strain, consistent with prior findings (33). Strikingly, PE_PGRS localization was highly dependent on culture conditions. When the bacteria were grown in the presence of Tween 80 detergent, which solubilizes much of the outer membrane (47), abundant PE_PGRS proteins were detected in the culture filtrate (CF) (**Fig. 3C**, lower panel). However, PE_PGRS proteins were entirely absent in the CF in the absence of Tween 80. In either condition, PE_PGRS proteins were found in the OBG fraction (**Fig. 3C**, upper panel).

### *PPE71* deletion upregulates stress response and carbon metabolism transcripts

Beyond a role in PE_PGRS secretion, the function of the *PPE71–38* locus remained unknown. To examine bacterial processes affected by *PPE71* gene dose, we subjected total bacterial RNA from WT+vector, WT+71OE, and Δ7138+vector strains to RNA-seq. Despite considerable within-group variability, principal component analysis demonstrated good separation of these strains by transcripts, particularly the WT+71OE strain (**Fig. 4A**). Functional annotation using the Database for Annotation, Visualization, and Integrated Discovery (DAVID) demonstrated a feature cluster for universal stress response genes that was significantly upregulated in the Δ7138+vector strain compared to WT+vector (**Fig. 4B**). Similar analysis identified a cluster of fatty acid biosynthesis genes significantly downregulated in the WT+71OE strain compared to WT+vector (**Fig. 4C**). While we noted differences in some stress response genes in the WT+71OE strain and fatty acid synthesis genes in the Δ7138 strain, neither of these clusters was significant for those comparisons. Notably, transcripts for *PE_PGRS* family genes did not vary consistently between the strains (**Table S1**).

**Figure 4:**
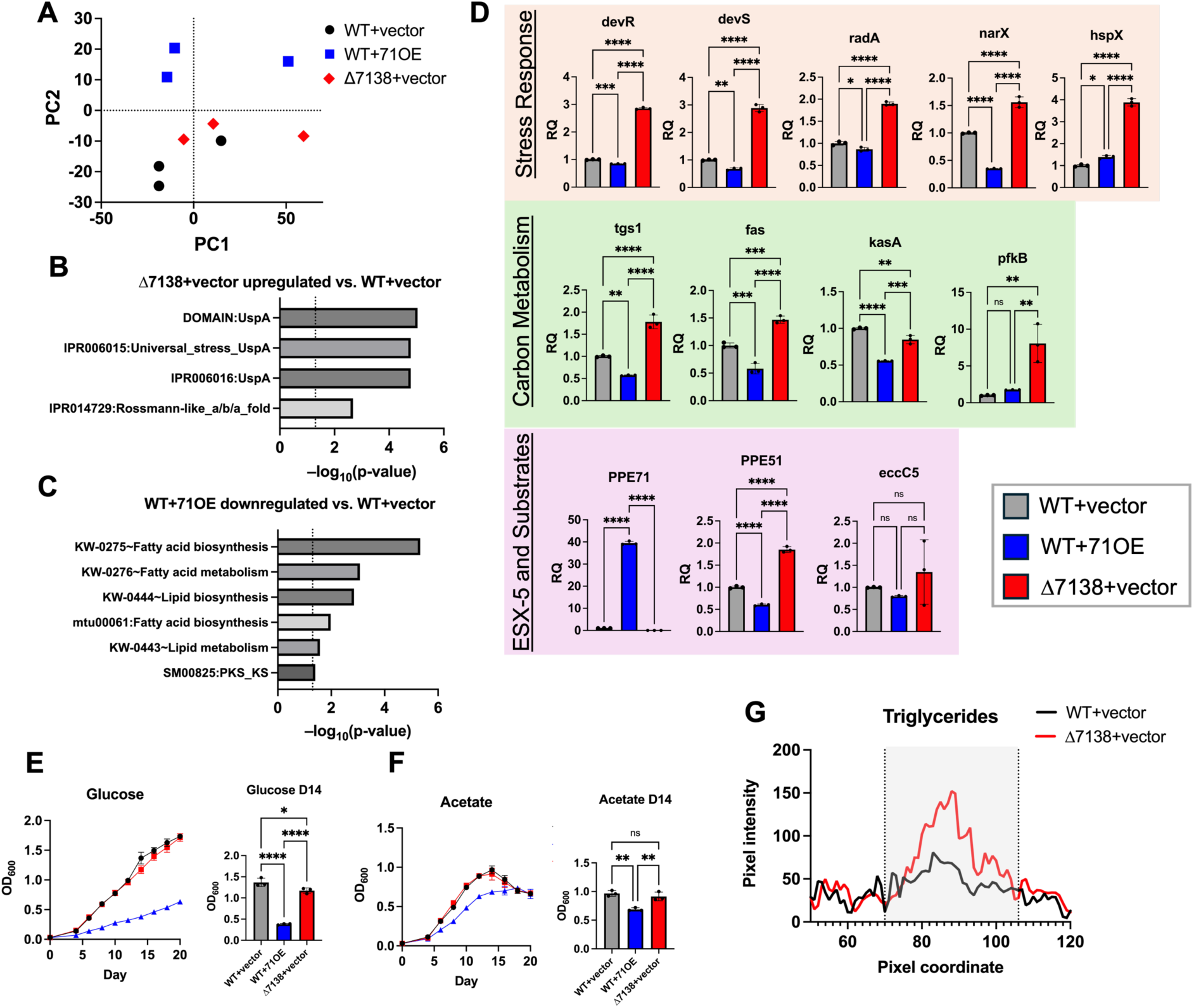
H37Rv Δ7138+vector upregulates stress response and fatty acid synthesis transcripts, while WT+71OE represses these genes. (A) Principal component plot of RNA-seq transcripts from WT+vector, WT+71OE, and Δ7138+vector strains (n=3). (B) Selected significant functional annotation terms for genes upregulated (1.5-fold change, p-adj < 0.05) in the Δ7138+vector strain compared to WT+vector. (C) Selected significant functional annotation terms for genes downregulated (1.5-fold change, p-adj < 0.05) in the WT+71OE strain compared to WT+vector. (D) Relative expression by qPCR of highly differentially expressed genes in WT+vector (grey), WT+71OE (blue) and Δ7138+vector (red) strains. Genes are grouped by function into stress response (orange), carbon metabolism (green) and ESX-5 and substrates (pink) pathways. For all genes, relative expression is normalized to the WT+vector strain. (n=3, ns: non-significant; *: p<0.05; **: p<0.01; ***: p<0.001; ****: p<0.0001 by one-way ANOVA.) (E–F) Growth curves of *M.tb* strains in minimal 7H9 broth supplemented with 0.05% tyloxapol and either (E) 0.2% glucose or (F) 0.2% acetate. Graphs of optical densities at 600 nm (OD_600_) for the Day 14 timepoints are provided for each medium. (n=3, ns: non-significant; *: p<0.05; **: p<0.01; ****: p<0.0001 by one-way ANOVA.) (G) Apolar lipids were extracted from H37Rv WT+vector and Δ7138+vector strains labeled with ^14^C-propionate, subjected to thin-layer chromatography (TLC) and visualized by phosphorimaging. Pixel intensities were measured across each lane of the TLC plate in the region surrounding the TG peak (origin = 0).

To validate the predicted functional annotations, we selected differentially expressed genes corresponding to stress response and carbon metabolism pathways, as well as several ESX-5-associated genes of interest, for qPCR studies. Among stress response genes, we found that *devR*, *devS*, *radA*, *narX*, and *hspX* were all significantly upregulated in the Δ7138+vector strain compared to WT+vector (**Fig. 4D**, top). These genes were also downregulated in the WT+71OE strain compared to WT, with the exception of *hspX*. Among carbon metabolism genes, *tgs1*, *fas*, and *pfkB* were upregulated in Δ7138+vector compared to WT+vector, while *tgs1*, *fas*, and *kasA* were downregulated in WT+71OE compared to WT+vector (**Fig. 4D**, middle). Hence, transcripts for the core components of fatty acid synthases I and II, as well as a triglyceride synthase, increase when *PPE71* is lost and decrease when it is overexpressed. To assess whether PPE71 expression influenced the ESX-5 secretion apparatus, we evaluated *eccC5* (a core ESX-5 component) mRNA levels. Expression of *eccC5* did not differ between the strains, suggesting that the presence or absence of *PPE71* does not affect ESX-5 expression (**Fig. 4D**, bottom). We observed differential expression of some *PE/PPE* family genes between strains; for example, *PPE51* was upregulated in Δ7138+vector and downregulated in WT+71OE compared to WT+vector. As a whole, the opposing effects seen in the Δ7138+vector strain and the WT+71OE strain support a role for *PPE71* gene dose on these stress response and lipid metabolism processes.

### WT+71OE has a carbon source-specific growth defect

Among the most differentially expressed genes between our strains were lipid synthesis genes and *pfkB*, an inducible phosphofructokinase that can act as the rate-limiting step in either glycolysis or gluconeogenesis (*48, 49*). Differential activity of these pathways may alter the ability of each strain to utilize particular carbon sources. As a result, we tested whether the Δ7138+vector or WT+71OE strains exhibited irregular growth behavior when provided with single carbon sources. When grown in minimal medium containing glucose as a carbon source, WT+71OE achieved an OD_600_ of only 0.38 by Day 14, compared to 1.4 for WT+vector (**Fig. 4E**). By contrast, WT+71OE achieved an OD_600_ of 0.67 (WT+vector: OD_600_ 0.97) by Day 14 in acetate-containing medium (**Fig. 4F**). Growth in glycerol-containing medium resembled that of the glucose condition (**Fig. S3A**). Propionate and propionamide permitted less growth but likewise demonstrated a growth defect specific to WT+71OE (**Fig. S3B–C**). Thus, while WT+71OE exhibited a general growth defect in the minimal media conditions, this effect was much more pronounced in the glucose medium. The Δ7138+vector strain showed few to no differences compared to WT+vector (**Figs. 4E–F, S3A–C**).

### Δ7138+vector accumulates more triglyceride than WT+vector

Because of the strong upregulation of fatty acid synthesis genes in the Δ7138+vector strain, we sought to determine whether this mutant showed any differences in the levels of different lipid species. We extracted apolar lipids from ^14^C-propionate-treated WT+vector and Δ7138+vector cultures and subjected them to thin-layer chromatography (TLC) (**Fig. S4A**). Upon quantifying the intensity of the peak corresponding to triglyceride (TG) lipids, we found that Δ7138+vector had higher levels of TGs than WT+vector (**Fig. 4G**). TG accumulation has been previously observed as a property of L2 strains, which frequently feature disruption of the *PPE71–38* locus (*50*). We do note, however, that the effect we observed is smaller than was previously reported for L2 strains (*50*).

### Increased *PPE71* expression increases the levels of PE/PPE proteins and lipoproteins in the *M.tb* cell wall

To more comprehensively examine changes in secreted proteins between the WT+vector, WT+71OE, and Δ7138+vector strains, we subjected CF and OBG fractions to protein mass spectrometry. Principal component analysis of CF (**Fig. 5A**) and OBG (**Fig. 5B**) fractions again demonstrated separation of the strains despite some within-group heterogeneity, particularly when grouped by OBG fraction. While only a few PE/PPE proteins were detected in the OBG fraction, these proteins exhibited an analogous pattern: PE25 and PPE19 were decreased in Δ7138+vector compared to WT+vector, while PE_PGRS38 was increased in WT+71OE compared to WT+vector (**Fig. 5C**). Hence, all three detected PE/PPE proteins were more abundant in the WT+71OE strain compared to the Δ7138+vector strain. Additionally, two PPE proteins as well as numerous Esp- and Esx-family type VII secretion system substrates were detected in the CF fraction (**Fig. S5**, **Table S2**).

**Figure 5:**
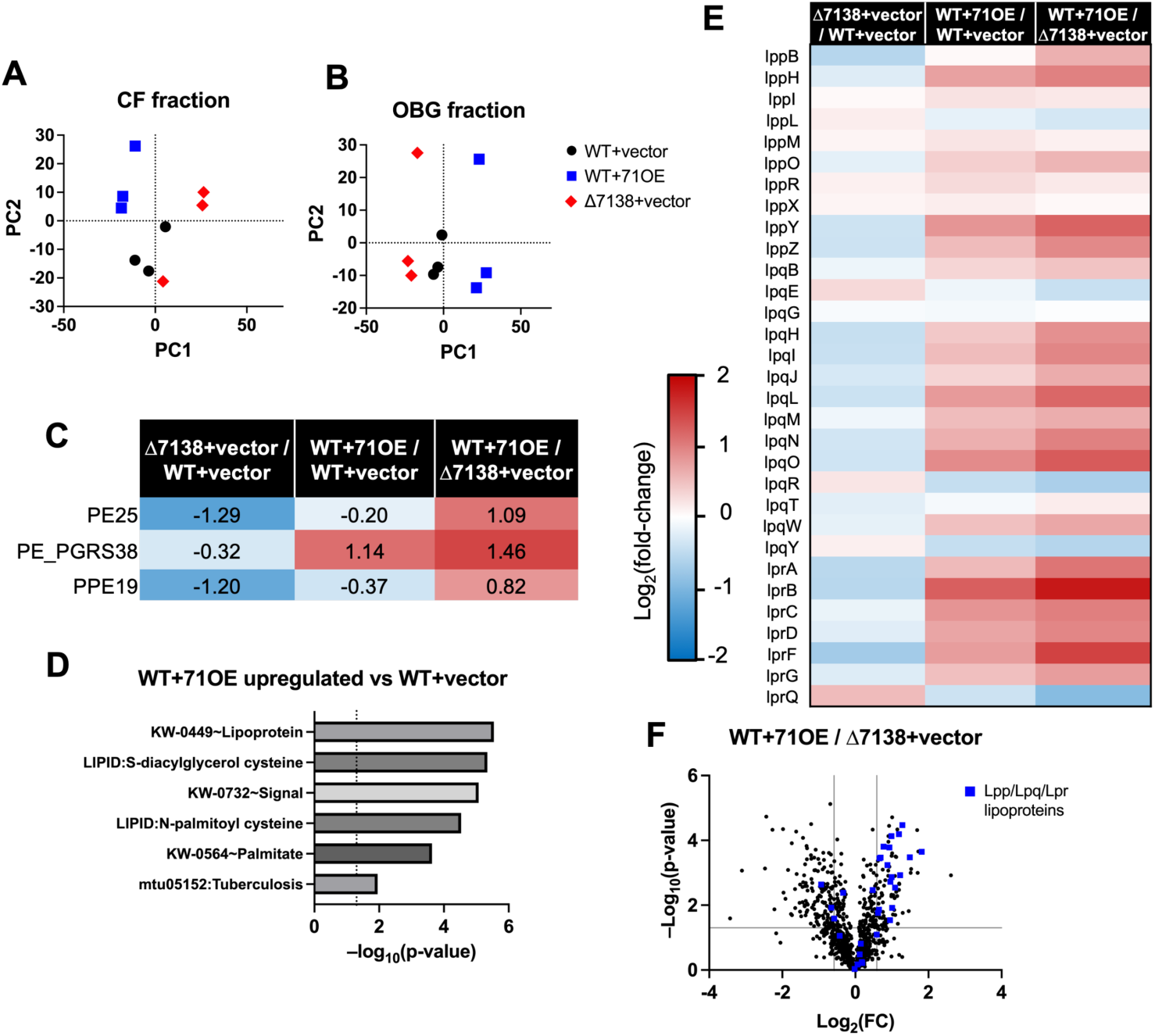
H37Rv WT+71OE upregulates lipoproteins in the outer mycomembrane. (A–B) Principal component plots of mass spectrometry readouts for culture filtrate (CF) (A) and octyl glucoside (OBG)-extracted (B) fractions from WT+vector, WT+71OE, and Δ7138+vector cultures (n=3). (C) Log_2_ fold-changes between each pair of strains for PE/PPE family proteins detected in OBG-extracted fractions by mass spectrometry. (D) Selected significant functional annotation terms for proteins upregulated (1.5-fold change, p < 0.05) in the OBG-extracted fraction of the WT+71OE strain compared to WT+vector. (E) Heat map of log_2_ fold-changes between each pair of strains for Lpp-, Lpq-, and Lpr-family lipoproteins detected in OBG-extracted fractions by mass spectrometry. (F) Volcano plot of proteins detected in OBG-extraction fractions for the WT+71OE vs Δ7138+vector comparison. The thresholds indicated are ± 1.5-fold change (vertical lines) and p < 0.05 (horizontal line). Lpp-, Lpq-, and Lpr-family lipoproteins are indicated with blue squares.

We performed functional annotation analysis on differentially abundant secreted protein levels using DAVID. Of particular note was a cluster associated with lipoproteins, which was upregulated in the OBG fraction of WT+71OE compared to WT+vector (**Fig. 5D**, **Table S3**). We generated a heat map including all 31 detected Lpp-, Lpq-, and Lpr-family lipoproteins and found that these proteins tended to be highly upregulated in the WT+71OE strain and subtly downregulated in the Δ7138+vector strain compared to WT+vector (**Fig. 5E**). Plotting these lipoprotein families on the overall volcano plot for the WT+71OE vs. Δ7138+vector comparison (OBG fraction) demonstrated that several were among the most differentially expressed genes and confirmed a pronounced directional shift in these families (**Fig. 5F**).

### Restoring *PPE71* expression in *M.tb* HN878 represses stress response and carbon metabolism transcripts

The L2 lineage strains of *M.tb* show a consistent phenotype marked by triglyceride accumulation and constitutive induction of DosR regulon targets (*50*). We were intrigued by the resemblance between this phenotype and the transcriptional changes we observed in the Δ7138+vector strain. In particular, we wondered if restoring expression of *PPE71* into *M.tb* HN878, a hypervirulent L2 strain with an IS*6110* insertion sequence disrupting all genes in the *PPE71–38* locus (*51*), would reverse this induction of stress response and carbon metabolism genes. We sequenced the *PPE71–38* locus of our HN878 isolate and confirmed the presence of an IS*6110* element that abolished the *esxX* and *esxY* genes as well as the *PPE71* gene C-terminus (after amino acid L26) and the *PPE38* gene N-terminus (before amino acid Y18) (**Fig. 6A**). These junctions are identical to a previously reported whole-genome sequence of *M.tb* HN878 (Genbank NZ_CM001043.1) (*51*). Next, we transformed an animal-passaged HN878 strain with the 71 and 71XY38 complementation constructs, the 71OE construct, and empty vector. By qPCR, we found that both native promoter-complemented strains (HN878+71 and HN878+71XY38) expressed around 1000-fold more *PPE71* transcripts than the HN878+vector strain, while HN878+71OE expressed around 15000-fold more *PPE71* than HN878+vector (**Fig. 6B**). These findings confirmed that HN878 does not natively express *PPE71* or *PPE38* and that the *PPE71* addback constructs were behaving as expected.

**Figure 6:**
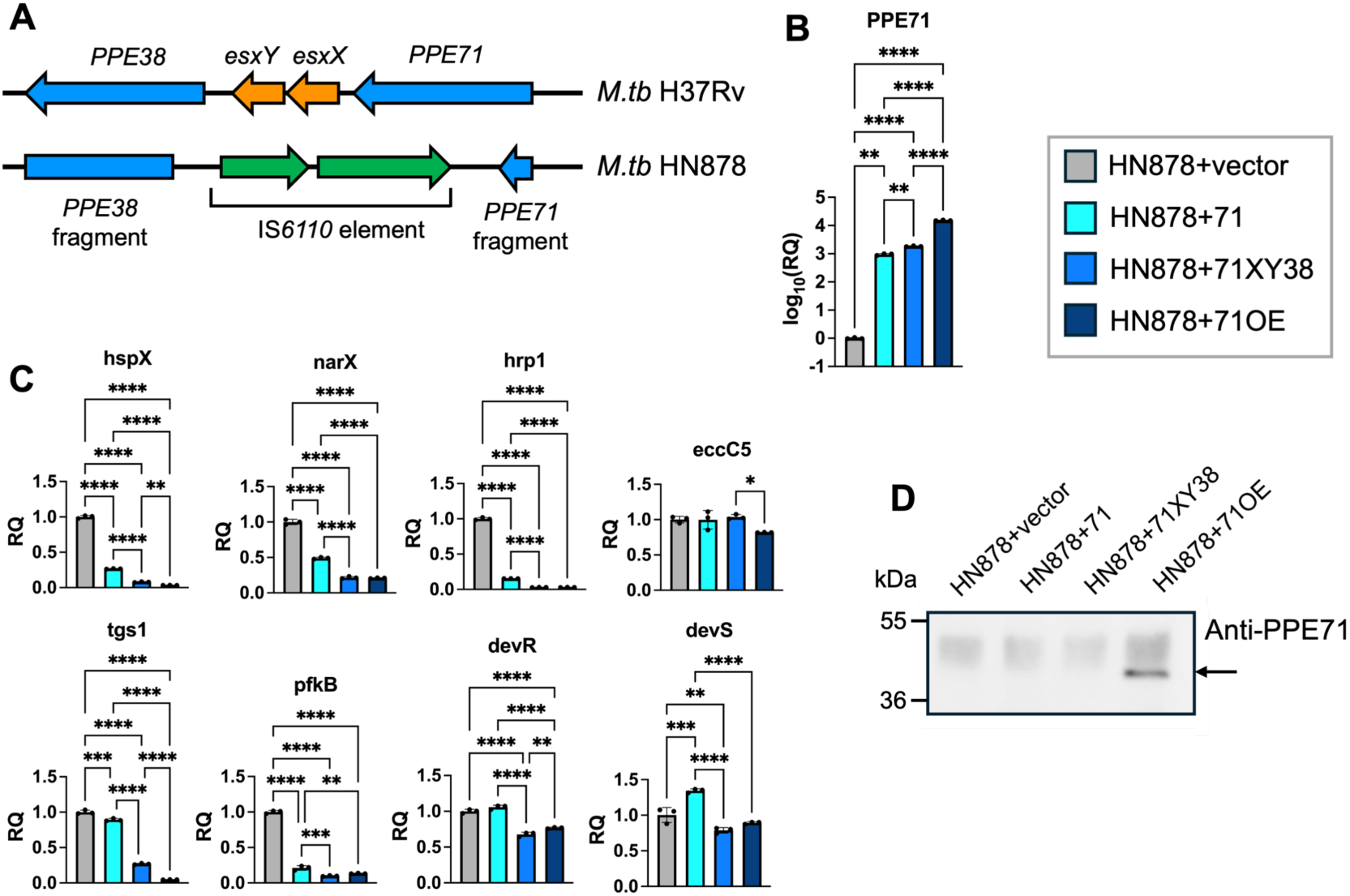
Re-expressing *PPE71* in the natural loss-of-function strain HN878 decreases stress response and carbon metabolism transcripts. (A) Comparison between the *PPE71–PPE38* loci in the *M.tb* strains H37Rv (L4) and HN878 (L2). The HN878 strain features an IS*6110* transposable element at this locus that disrupts the C-terminus of *PPE71* (including almost all of the gene), *esxX*, *esxY*, and the N-terminus of *PPE38* (including the start codon). The *PPE71* fragment in *M.tb* HN878 includes amino acids M1–L26, while the *PPE38* fragment includes Y18–*392. (B) Relative expression (log_10_ fold-change) by qPCR of *PPE71* in the *M.tb* HN878 addback strains, normalized to baseline expression in the HN878+vector strain. *M.tb* 16S rRNA was used as the internal control for normalization. (mean±SD; n=3, **: p<0.01; ****: p<0.0001 by one-way ANOVA.) (C) Relative expression by qPCR of select stress response and carbon metabolism genes, with eccC5 as a control. *M.tb* 16S rRNA was used as the internal control for normalization. All significant comparisons by one-way ANOVA are depicted. (mean±SD; n=3, *: p<0.05; **: p<0.01; ***: p<0.001; ****: p<0.0001.) (D) Western blot of whole-cell lysate from HN878 *PPE71* variant strains blotted with anti-PPE71 antibody. PPE71 protein is visible in the HN878+71OE strain and is indicated with an arrow. (kDa: kilodaltons.)

Subsequently, we tested a selection of stress response and metabolic transcripts that were dysregulated in the H37Rv Δ7138+vector strain relative to the WT+vector strain. We found that all of *hspX*, *narX*, *hrp1*, *tgs1*, and *pfkB* had significantly decreased expression with *PPE71* addback into HN878 (**Fig. 6C**). Excitingly, the repression of these genes by *PPE71* addback was dose-dependent: the HN878+71XY38 strain (one copy of PPE71 and one copy of its close paralogue PPE38) exhibited a greater decrease than the HN878+71 strain (one copy of PPE71), while the HN878+71OE strain showed the strongest decrease. Upon examining *devR* and *devS* transcripts, we found smaller but significant decreases in the HN878+71XY38 and HN878+71OE strains but not the HN878+71comp strain compared to WT. As a control, the level of *eccC5* transcript showed little change aside from a decrease in the HN878+71OE strain.

Additionally, we extracted ^14^C-propionate-labeled apolar lipids from these HN878 *PPE71* variant strains. However, we did not observe any differences in lipids between these strains (**Fig. S4B**). Upon blotting lysates from the HN878 *PPE71* variant strains with the antibody against PPE71, we observed a band consistent with PPE71 protein (∼37 kDa) in the HN878+71OE strain (**Fig. 6D**). This is an analogous pattern to the H37Rv *PPE71* variant strains, in which the antibody detected overexpressed levels of *PPE71*. Additionally, we blotted the OBG fractions of the HN878 *PPE71* variant strains with anti-PE_PGRS33 and observed a broader range of PE_PGRS family proteins in the HN878+71XY38 and HN878+71OE strains compared to either HN878+vector or HN878+71 (**Fig. S6**). Hence, restoration of the *PPE71* locus or *PPE71* overexpression appeared to increase PE_PGRS protein secretion in *M.tb* HN878.

## Discussion

The widespread loss-of-function polymorphisms at the *M.tb PPE71–38* locus suggest that this genetic disruption benefits the pathogen. Here, we present a mechanism by which *PPE71–38* deletion may enhance *M.tb* virulence. Deletion of *PPE71–38* in the *M.tb* H37Rv background led to upregulation of stress response transcripts, including the *devR/devS* genes that control adaptation to dormancy, and fatty acid synthesis genes. Conversely, *PPE71* overexpression led to downregulation of these same genes, supporting the causal role of *PPE71* dose on these processes. Overexpressing *PPE71* also broadly increased the abundance of lipoproteins in the mycobacterial cell wall, which may reflect underlying changes in lipid metabolism and cell wall processes. Lastly, restoring *PPE71* expression in *M.tb* HN878, a hypervirulent L2 strain with a disrupted *PPE71* locus, demonstrated that *PPE71* is sufficient to repress these stress response and metabolic pathways.

Vital prior work in the field has established that PPE71 is necessary for the secretion of PE_PGRS family proteins in mycobacteria (33). Past proteomic studies demonstrated a broad loss of PE_PGRS proteins in the culture filtrate of *M.tb* strains upon PPE71–38 deletion (33, 34). Here we have demonstrated that increasing *PPE71* gene dose to far above WT levels can greatly increase PE_PGRS export. Since PPE71–38 and several PE_PGRS proteins are among the most abundant *M.tb* proteins detected in animal infection, it is likely that *M.tb* induces *PPE71–38* and thus PE_PGRS secretion *in vivo* by natural processes (38). As we did not detect appreciable PPE71–38 protein levels in WT H37Rv, it is also possible that this strain exhibits less expression of *PPE71–38* than has been previously noted in *M.tb* CDC1551 (33). Hence, *PPE71–38* expression and thus PE_PGRS protein levels may vary even between strains with intact *PPE71– 38* loci. We determined that *M.tb* PPE71 localizes to the outer mycomembrane and is cell surface exposed, which is also the case for the *M. marinum* homolog of *PPE38* (42). In addition, we have found that the native compartment for PE_PGRS proteins is likely the outer mycomembrane rather than the soluble culture filtrate. This is consistent with the recently proposed ‘sail’ structure for PGRS domains that would anchor them in the outer leaflet of the mycomembrane, as well as a recent characterization of the *M.tb* outer membrane proteome (*52, 53*). While it remains possible that some PE_PGRS proteins may be solubly secreted or may detach from the mycomembrane during infection, our findings suggest that family is predominantly borne on the outer mycomembrane. This dramatic increase in PE_PGRS protein secretion was a major factor in our decision to pursue the 71OE construct in this study. However, our 71OE construct uses a non-integrating plasmid, and the gradual loss of this construct during the late animal timepoints may explain why the Δ7138+71OE strain appears to be attenuated at Week 4 but lose this attenuation at Week 8. Since this limitation does not arise until after Week 4, it is unlikely to be present in our shorter duration *in vitro* experiments. Additionally, it is worth considering that constitutive overexpression of a single gene, as in the case of the 71OE strains, may introduce additional artificial effects beyond the role of *PPE71* itself.

It is initially unintuitive why loss of *PPE71–38* and thus PE_PGRS secretion would be tolerated by *M.tb*, much less under apparent positive selection across diverse clinical isolates. Here, we explain this observation by highlighting beneficial effects on stress response and lipid metabolism in our Δ7138+vector strain. *M.tb* lipid metabolism is intricately involved in virulence, and mycobacteria enriched in lipids show improved resistance to stressful conditions and tolerance to certain antibiotics (*54–57*). Several of the stress response pathways we highlight, including the *devR* operon, regulate the same metabolic genes that we see as differentially expressed, most prominently *pfkB* and *tgs1* (*58, 59*). Hence, it is possible that the upregulated stress response genes we observe in the Δ7138 mutant and the L2 strain HN878 may be driving the metabolic effects here. Clinical L2 strains of *M.tb* constitutively overproduce triglycerides, and we observed a similar phenotype in our H37Rv Δ7138+vector strain as well, although our finding was noticeably smaller (*50*). Intriguingly, restoration of *PPE71* full locus expression or *PPE71* overexpression into HN878 had a comparatively weaker effect on direct repression of *devR* and *devS* transcripts compared to the other transcripts we measured. L2 strains are known to have a mutation in the *dosT* sensor as well as single nucleotide polymorphism in the *devR* promoter, both of which drive regulon expression (*60, 61*). The presence of multiple co-occurring genetic lesions in L2 strains that each boost the DosR regulon establishes the importance of this regulon for the L2 phenotype and may explain why adding back *PPE71* did not suppress the *devR/S* genes to the same extent seen with other hits. Hence, we believe that *PPE71* has a causal role in regulating mycobacterial stress response and metabolic pathways and that this role may be partly mediated through the DosR system. Of course, it remains possible that loss of PE_PGRS or PPE-MPTR proteins in the mycomembrane may directly or indirectly mediate these effects. Additionally, L2 strains feature a variety of polymorphisms at the *PPE71– 38* locus, including various deletions, insertion sequences at different locations and orientations, and chimeric *PPE71-38* gene fusions (*30, 31*). These diverse mutations may impair the function of the locus to differing extents. It is unknown, for example, whether a *PPE71*-*38* chimera (accompanied by a loss of *esxX* and *esxY*) remains functional, although our finding that the Δ7138+71 strain did not suppress hypervirulence *in vitro* suggests that a single *PPE71* copy under its native promoter may be insufficient to restore proper function.

The chemical and material composition of the mycomembrane must be carefully controlled to permit uptake of necessary materials while limiting exposure to toxic host-derived species. By adjusting the levels of PPE71 and thus PE_PGRS proteins anchored into the mycomembrane, *M.tb* may modulate key properties of the outer mycomembrane. As an intriguing example, we found that the expression of *PPE51* tracked inversely with that of *PPE71*. PPE51 has been implicated in the uptake of small carbon sources, such as glucose, glycerol, and propionamide (*62*). Consistent with this, the WT+71OE strain exhibited impaired growth in glucose but not acetate. Additionally, we observed a striking increase in over three quarters of detected Lpp, Lpq, and Lpr lipoproteins in the cell wall of the WT+71OE strain. While these proteins serve diverse functions, many of their known activities have been linked to the mycomembrane itself, such as nutrient acquisition and maintaining cell wall integrity (*63, 64*). These results collectively support a model in which the consequences of changing *PPE71* dosage may occur downstream of changes in outer mycomembrane composition or permeability to small molecules. Future work would be required to examine any differences in small molecule uptake and cytoplasmic metabolite levels in strains with diverse *PPE71* expression, including how these differences may impact the activities of antitubercular drugs.

In sum, our findings provide insight into the broad distribution of *PPE71–38* locus loss-of-function polymorphisms across *M.tb* strains. Per our results, disruption of *PPE71–38* upregulates stress response and lipid synthesis transcripts, while *PPE71* overexpression or restoration of *PPE71–38* expression into a natural deletion strain represses these same processes. We observed broad shifts in the levels of PE_PGRS and lipoproteins in the cell wall due to changes in *PPE71* expression as well as changes in carbon source preference and triglyceride production. Given the increasing global concern of hypervirulent *M.tb* L2 strains with *PPE71–38* polymorphisms, further exploration of these interlinked processes may uncover regulatory steps vital to *M.tb* pathogenesis and tuberculosis intervention.

## Materials and methods

### Bacterial strains and media

*M.tb* strains were cultured in Middlebrook 7H9 broth supplemented with 0.5% glycerol, 10% OADC, and 0.05% Tween 80 (henceforth ‘complete 7H9’) or on Middlebrook 7H11 agar supplemented with 0.5% glycerol and 10% OADC. Single carbon source media were prepared by supplementing minimal 7H9 broth containing 0.05% tyloxapol with each of 0.2% glucose, 0.2% glycerol, 0.2% acetate, 0.1% propionate, or 5 mM propionamide and adjusting to pH 7.2, as described in previous work (*62, 65*). Propionate/MES induction medium (henceforth ‘induction medium’) was prepared by supplementing 7H9 broth with 0.1% glycerol, 1 mM sodium propionate, and 100 mM 2-(4-morpholino)-ethane sulfonic acid (MES) and adjusting to pH 6.5, adapted from prior work (*66*). For lipid extraction, strains were cultured in modified 7H9 broth supplemented with 0.5% glycerol, 10% OADC, 0.1 mM sodium propionate and 0.05% Tyloxapol, adapted from prior work (*67*). Cultures were grown at 37°C with shaking at 180–200 rpm. Growth curves were conducted by seeding from mid-log phase cultures to the equivalent of OD_600_ 0.03 in 10 mL. Selection was achieved using 25 ug/mL kanamycin or 50 ug/mL hygromycin as needed.

For plasmid cloning, *E.coli* DH5*α* was grown in LB broth or on LB agar at 37°C. For recombinant PPE71 protein production, *E.coli* BL21 was grown analogously. Selection was achieved using 100 ug/mL carbenicillin or 50 ug/mL kanamycin as needed.

### *M.tb* mutant construction

The *PPE71–38* locus consists of four genes: *PPE71*, *esxX*, *esxY*, and *PPE38*. *PPE71* is identical to *PPE38* (Rv2352c) with the exception of a 7 amino acid deletion (GGAGAGM) spanning amino acids 357–363 of the 391-residue PPE38 protein (30). Hence, we believe these two proteins are likely functionally redundant. Due to a later-noted error in the assembly of the H37Rv genome (*30, 68*), *PPE71*, *esxX*, and *esxY* each lack Rv numbers. These genes can instead be identified by their MT numbers: MT2422 (*PPE71*), MT2421 (*esxX*), MT2420 (*esxY*), and MT2419 (*PPE38*). We confirmed by sequencing that all four genes are present in the *M.tb* H37Rv strain used in this work. Indeed, the *PPE71–38* locus of our *M.tb* H37Rv isolate is identical to that of the CDC1551 NC_002755.2 complete sequence, which properly includes all four genes in the locus (*69*).

*M.tb* H37Rv Δ*PPE71-esxX-esxY-PPE38* (henceforth Δ7138) was generated using a two-step specialized transduction design, described in prior work (*70*). In brief, a sequence-specific phage was used to replace the region between Tyr18 of PPE71 and Ala373 of PPE38 with a *sacB*-*hyg* cassette, and the transformant was selected on 7H11 agar supplemented with 50 ug/mL hygromycin. Next, a resolving phage was used to excise the cassette to leave an unmarked deletion, and the transformant was counter-selected on 7H11 agar supplemented with 10% sucrose. Deletion was confirmed by PCR amplification with dPPE38_57_f/r followed by Sanger sequencing with dPPE38_inner_f/r. Note that, although the most widely available reference sequence for *M.tb* H37Rv depicts the *PPE71–PPE38* locus as abridged to contain only *PPE38*, in actuality *M.tb* H37Rv has an intact *PPE71–PPE38* locus including all four of *PPE71*, *esxX*, *esxY*, and *PPE38*. This error in the H37Rv reference sequence assembly was noted previously (*30, 68*).

To construct the *PPE71* single gene complement, plasmid pMH94 (*71*) was digested with XbaI. The PPE71 fragment was amplified from *M.tb* H37Rv genomic DNA with PPE71_24f/r and incorporated into the linearized plasmid by Gibson assembly to produce pMH94-PPE71. Sequences were verified using pMH94_Fseq/Rseq.

To construct the *PPE71-esxX-esxY* complement (henceforth 71XY) and the *PPE71-esxX-esxY-PPE38* (henceforth 71XY38) complement, plasmid pMH94 was digested with KpnI and XbaI. The 71XY fragment was amplified from *M.tb* H37Rv genomic DNA with PPE71locus_59_F and EsxY_59_R. The 71XY38 fragment was amplified from *M.tb* H37Rv genomic DNA with PPE71locus_59_F and PPE38term_59_R. Each fragment was incorporated into the linearized plasmid using cut-and-paste ligation following digestion with KpnI and XbaI to produce pMH94-71XY and pMH94-71XY38. Sequences were verified using pMH94_Fseq/Rseq as well as PPE71-EsxX_Fseq/Rseq and EsxY-PPE38_Fseq/Rseq for internal coverage of the large fragments.

To construct the *PPE71* overexpressor (henceforth 71OE), plasmid pSD5 (*72*) was digested with NdeI and MluI. The PPE71 fragment was amplified from *M.tb* H37Rv genomic DNA with PPE71_pSD5_f/r and incorporated into the linearized plasmid by Gibson assembly to produce pSD5-71OE. Sequences were verified using pSD5_insert_FWD/REV.

*M.tb* H37Rv (WT) was electroporated with pMH94 and pSD5-71OE to produce WT+vector and WT+71OE strains, respectively. *M.tb* H37Rv Δ7138 was electroporated with pMH94, pMH94-PPE71, pMH94-71XY, pMH94-71XY38, and pSD5-71OE to produce Δ7138+vector, Δ7138+71, Δ7138+71XY, Δ7138+71XY38, and Δ7138+71OE, respectively. *M.tb* HN878 was electroporated with pMH94, pMH94-PPE71, pMH94-71XY38, and pSD5-71OE to produce HN878+vector, HN878+71, HN878+71XY38, and HN878+71OE, respectively.

Protocols for *M.tb* genomic DNA isolation and electroporation of foreign DNA into *M.tb* have been previously described (*73*). Plasmids and oligonucleotides used in this work are provided in **Tables S4** and **S5**, respectively.

### RNA isolation

*M.tb* strains were grown in 50 mL cultures to mid-log phase. Bacterial pellets were washed and transferred to 50 mL induction medium, in which they were cultured for 3 days at 37°C with shaking. Total *M.tb* RNA isolation was performed as described previously (*74*). Briefly, *M.tb* cultures were lysed in 1 mL TRIzol (Thermo) by bead beating in a Precellys Evolution homogenizer for 3 cycles of 7400 bpm for 30 sec, with chilling on ice between cycles. Samples were subjected to phenol-chloroform extraction followed by two incubations with DNase I (Qiagen) to remove residual DNA. Samples were then purified using an RNeasy kit (Qiagen) per manufacturer’s instructions. RNA concentration and purity was measured by 260/280 ratio on a NanoDrop spectrophotometer (Thermo).

### Real-time quantitative PCR

Purified *M.tb* RNA was converted to complementary DNA (cDNA) using an iScript cDNA Synthesis kit (Bio-Rad), per manufacturer instructions. Quantitative PCR (qPCR) was performed using iTaq Universal SYBR Green Supermix (Bio-Rad), per manufacturer instructions, on a QuantStudio 3 Real-Time PCR System (Applied Biosystems) using a ΔΔCt method. Transcript levels were normalized to expression of *M.tb* 16S rRNA. Primer pairs for genes targeted in this work are provided in **Table S5**.

### RNA-seq

Purified *M.tb* RNA was subjected to DNA and RNA Qubit fluorimetry to verify the absence of DNA contamination and precisely quantify RNA amounts. RNA samples were depleted of rRNA and converted to cDNA. Samples were subjected to Illumina next-generation sequencing, with 30 million 2×150bp paired-end reads obtained per sample. Reads were trimmed to remove adapter sequences and mapped to the *M.tb* H37Rv genome using the Bowtie2 tool (*75*).

Differential gene expression between each pairwise comparison of strains was conducted using the Wald test in the DESeq2 tool (*76*) corrected using a Benjamini-Hochberg multiple test correction. Functional annotation analysis was conducted by inputting a list of differentially expressed genes (≥1.5-fold differential expression, p-adjusted < 0.05) into the DAVID informatics database (*77, 78*).

### PPE71 protein production

To construct the pET28a-PPE71 plasmid for inducible expression of PPE71 protein in E.coli, plasmid pET-28a(+) (Novagen) was digested with NdeI and SalI. The *PPE71* gene was amplified from *M.tb* H37Rv genomic DNA using PPE71_34F/R and incorporated into the linearized plasmid by Gibson assembly to produce pET28a-PPE71. Sequences were verified using T7_FWDseq/REVseq.

The pET28a-PPE71 expression construct was transformed into *E.coli* BL21 by heat shock. A successful transformant line was grown overnight in 100 mL of LB at 37°C, then 1 mM IPTG (Sigma) was added, and the culture was allowed to induce at 18°C for 24 hr. Cells were lysed by sonication on ice, and inclusion bodies were washed and denatured as described previously (*79*). PPE71 was purified using the in-frame 6xHis tags via a Cytiva HisTrap HP 5mL column per manufacturer instructions.

### *M.tb* protein extraction

To obtain *M.tb* whole-cell lysate (WCL), 10 mL mid-log phase bacterial cultures grown in complete 7H9 were lysed in 500 uL Tris-buffered saline (TBS) by bead beating in a Precellys Evolution homogenizer for 3 cycles of 7400 bpm for 30 sec, with chilling on ice between cycles. Protease Inhibitor Cocktail for bacterial cell extracts (Sigma) and 100 mM phenylmethylsulfonyl fluoride (PMSF) (Sigma) were each added at 1:100 dilution prior to lysis and are henceforth referred to as ‘protease inhibitors.’ Samples were pelleted, and the soluble fraction was passed through an 0.22 um cellulose acetate filter (Corning) to remove debris. For the proteinase K assay, bacteria were incubated with 2 ug/mL proteinase K (New England Biolabs) for 0, 5, 10, or 15 min at 37°C prior to lysis, as adapted from prior work (*42, 80*). Proteinase K activity was quenched by addition of protease inhibitors followed by immediate heat inactivation at 95°C.

To obtain *M.tb* culture filtrate (CF) and octyl glucoside (OBG) fractions, 20 mL bacterial cultures were grown for 3 days in a modified complete 7H9 medium containing no Tween 80. Each culture was split in half, Tween 80 was added to one half at a final concentration of 0.05%, and cultures were allowed to grow for an additional 24 hr at 37°C with shaking. Bacteria were pelleted, and the soluble CF fraction was subjected to trichloroacetic acid (TCA) precipitation, as described previously (*81*). Bacterial pellets were washed twice with TBS and incubated in 1% OBG (Sigma) in TBS with protease inhibitors for 30 min at 37°C to extract the outer mycomembrane fraction. Following incubation with OBG, the bacteria were pelleted, and the detergent-solubilized fraction was taken as the OBG fraction.

### Western blotting

Samples were boiled at 95°C for 10 min in 4X Laemmli sample buffer (Bio-Rad) and loaded on 4-15% Mini-PROTEAN TGX Gels (Bio-Rad) for electrophoresis at 100 V for approximately 1 hr. Proteins were transferred to 0.22 um PVDF membranes (Bio-Rad) by wet transfer method at 80 V for 45 min on ice. Membranes were blocked for 1 hr in 5% milk blocking buffer in TBS-T (TBS with 0.1% Tween 20) at room temperature. Primary antibodies were incubated overnight at 4°C in 5% milk, while secondary antibodies were incubated for 1 hr at room temperature in TBS-T. Membranes were exposed using SuperSignal West Pico PLUS substrate (Thermo) and imaged with a Kwik Quant imager (Kindle Biosciences). Protein sizes were estimated using the PageRuler Plus Prestained Protein Ladder (Thermo).

Anti-PPE71 (rabbit polyclonal) was obtained by immunization with the Gly362–Ser376 predicted epitope at the C-terminus of PPE71 (Sino Biological). Anti-FtsZ (rabbit antiserum) was obtained from BEI Resources, NIAID, NIH: Polyclonal Anti-Mycobacterium tuberculosis FtsZ (Gene Rv2150c) (antiserum, Rabbit), NR-44103. Anti-PE_PGRS33 (mouse monoclonal) 7C4.1F7 was obtained from the International AIDS Vaccine Initiative and was originally developed in previous work (*44*). Anti-rabbit and anti-mouse secondary antibodies were purchased from Cell Signaling Technologies (#7074 and #7076, respectively).

### Mass spectrometry

To obtain CF and OBG samples for proteomic analysis, 50 mL *M.tb* cultures were grown to mid-log phase, then induced in induction medium for 3 days. CF and OBG fractions were prepared as described above, except that CF fractions were first concentrated using 3 kilodalton (kDa) molecular weight cut-off columns (Cytiva) at 4°C, with added protease inhibitors. Samples were reduced with tris(2-carboxyethyl)phosphine (TCEP), alkylated with iodoacetamide, and further reduced with dithiothreitol (DTT). Samples were digested with endoprotease Lys-C and trypsin before being labeled with TMTpro 18-plex reagents (Thermo). Labeled peptide samples were pooled and analyzed on an Orbitrap Eclipse mass spectrometer (Thermo) with a FAIMS device. MS2 spectra were searched against an *M.tb* composite database using the COMET tool, and peptide-spectrum matches were filtered to a false-discovery rate of 1% (*82*). Peptide abundances were normalized to the total peptide abundance for each sample. Functional analysis was conducted as in the RNA-seq workflow.

### *M.tb* lipid extraction

To obtain *M.tb* lipids, 20 mL *M.tb* cultures were grown in complete 7H9 to OD_600_ 1.5. At 24 hr prior to harvest, 1 μCi/mL ^14^C-propionate was added to each culture. Lipid extraction was performed as adapted from prior work (*83*). Briefly, cell pellets were washed with phosphate-buffered saline (PBS) and incubated for 24 hr in 1:2 chloroform:methanol followed by a 48 hr incubation in 2:1 chloroform:methanol. Organic fractions were washed twice with distilled water, then dried under airflow.

### Thin-layer chromatography

In preparation for TLC, extracted lipid samples were resuspended in a volume of dichloromethane normalized by sample mass to equalize the concentration of each sample. Uptake of radiolabel was measured via scintillation count on a Beckman LS 6000SE, and 100,000 counts per minute (cpm) of each sample was loaded onto aluminum-backed silica gel TLC plates and allowed to run in one dimension using 98:2 petroleum ether:acetone, as described previously (*83*). TLC plates were exposed to a phosphorimaging cassette for 24 hours before being visualized on an Amersham Typhoon RGB phosphorimager. Background correction and semiquantitative band detection was conducted using ImageQuant TL software. The plot profile feature in ImageJ was used to measure pixel intensities across each TLC lane. Mean background pixel intensities were subtracted as determined for each image.

### Mouse infection and endpoints

Eight-week-old female BALB/cJ mice (#000651) were purchased from The Jackson Laboratory. Mice were housed within an animal biosafety level 3 facility and provided *ad libitum* rodent chow and clean water. The facility was maintained on alternating 12-hour light/dark cycles. Mice were infected with the *M.tb* strains described above by aerosol route using a Glas-Col Inhalation Exposure System with an inoculum of approximately 400 CFU per animal. To assess inocula, animals were sacrificed at Day 1 (n=3–4 per group), and homogenate from the combined left and right lungs were plated. At Week 4 and Week 8 timepoints, 8 animals were sacrificed per group. Right lungs and spleens were subjected to bead beating in a Precellys Evolution homogenizer and plated on 7H11 plates as a 10-fold dilution series to determine organ CFUs. Left lungs were fixed in 10% neutral-buffered formalin (NBF) for 2 days and subjected to histological embedding and sectioning, followed by staining with hematoxylin and eosin (H&E) to visualize tissue morphology. To determine total lung area and lung lesion area, H&E images were color-thresholded in ImageJ to quantify tissue between 0–230 brightness (total area) and 0–210 (lesion area), followed by outlier removal at a 5 pixel radius.

### Statistical analysis

Statical analysis was performed in GraphPad Prism Version 10. Groups were analyzed for significant findings using one-way analysis of variance (ANOVA) tests followed by Tukey’s test for multiple comparisons. Error bars in all figures represent mean ± standard deviation. All raw points are shown, and no points were omitted. All data points represent distinct samples; samples were not measured repeatedly. In all cases, *p* < 0.05 or a more stringent cutoff was used as a threshold for statistical significance.

### Ethics statement

Experimental procedures were conducted in accordance with protocol #MO22M466, approved by the Institutional Animal Care and Use Committee (IACUC) of Johns Hopkins University. No animals reached mandatory endpoints for humane euthanasia during the experiment, and no distress or pathology was noted in any animals throughout the experiment.

## Acknowledgements

We thank Dr. Eric Nuermberger for providing the HN878 strain used in this study. We also thank the members of the Thermo Fisher Scientific Center for Multiplexed Proteomics at Harvard Medical School for conducting the mass spectrometry quantification.

## Author contributions

Conceptualization: BK, WRB

Methodology: BK, CS, SR, WRJ, WRB

Investigation: BK, CS, SR, SS, YM-M, MG, JS

Supervision: WRB

Writing: BK, SS, WRJ, WRB

## Data and materials availability

All data are available in the main text or the supplementary materials. Raw transcriptomic data have been provided as a Microsoft Excel file, **Table S1**. Raw proteomic data for CF and OBG fractions have been provided as Microsoft Excel files, **Table S2** and **Table S3**, respectively. Plasmids or strains used in this study are available upon request to the corresponding author.

