## Supplemental Figures for "Loss of the *PPE71-esxX-esxY-PPE38* locus drives adaptive transcriptional responses and hypervirulence of *Mycobacterium tuberculosis* Lineage 2"

Benjamin Koleske *et al.*

**This PDF file includes:**

Figures S1 to S6
Tables S1 to S5

N.B.: Due to their large size, Tables S1 to S3 are each attached as separate Microsoft Excel files.

### Supplementary Figures

**A**

| Strain | Day 1 $\log_{10}$(CFU) |
|---|---|
| WT+vector | 2.55 |
| WT+71OE | 2.39 |
| Δ7138+vector | 2.65 |
| Δ7138+71 | 2.75 |
| Δ7138+71XY | 2.61 |
| Δ7138+71XY38 | 2.80 |
| Δ7138+71OE | 2.78 |

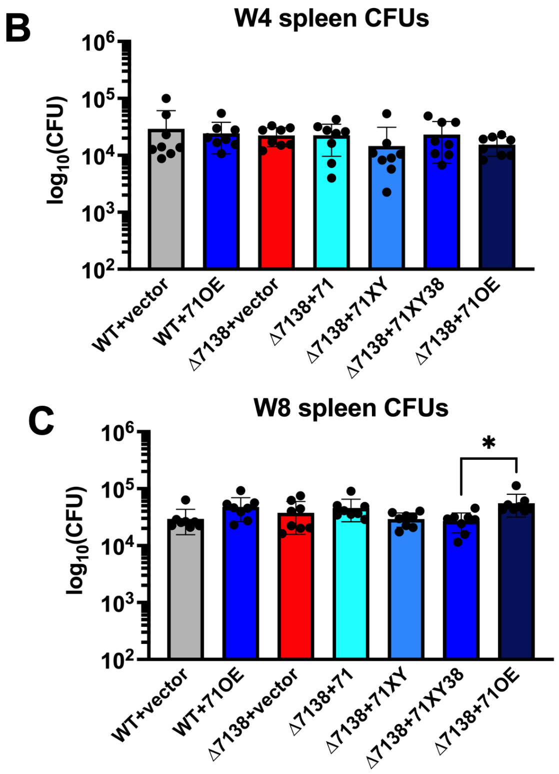


**Figure S1: H37Rv *PPE71* variant strains showed equivalent inocula and minimal spleen findings.**
(A) Day 1 lung CFUs to assess lung inocula for BALB/c infection with *M.tb* H37Rv *PPE71* variant strains. No strains had significantly different inocula from the WT+vector strain by one-way ANOVA. (n=3-4.)
(B–C) Spleen CFUs from Week 4 (E) and Week 8 (F) timepoints. All significant comparisons by one-way ANOVA are depicted. (mean±SD; n=8 each, *: $p<0.05$.)

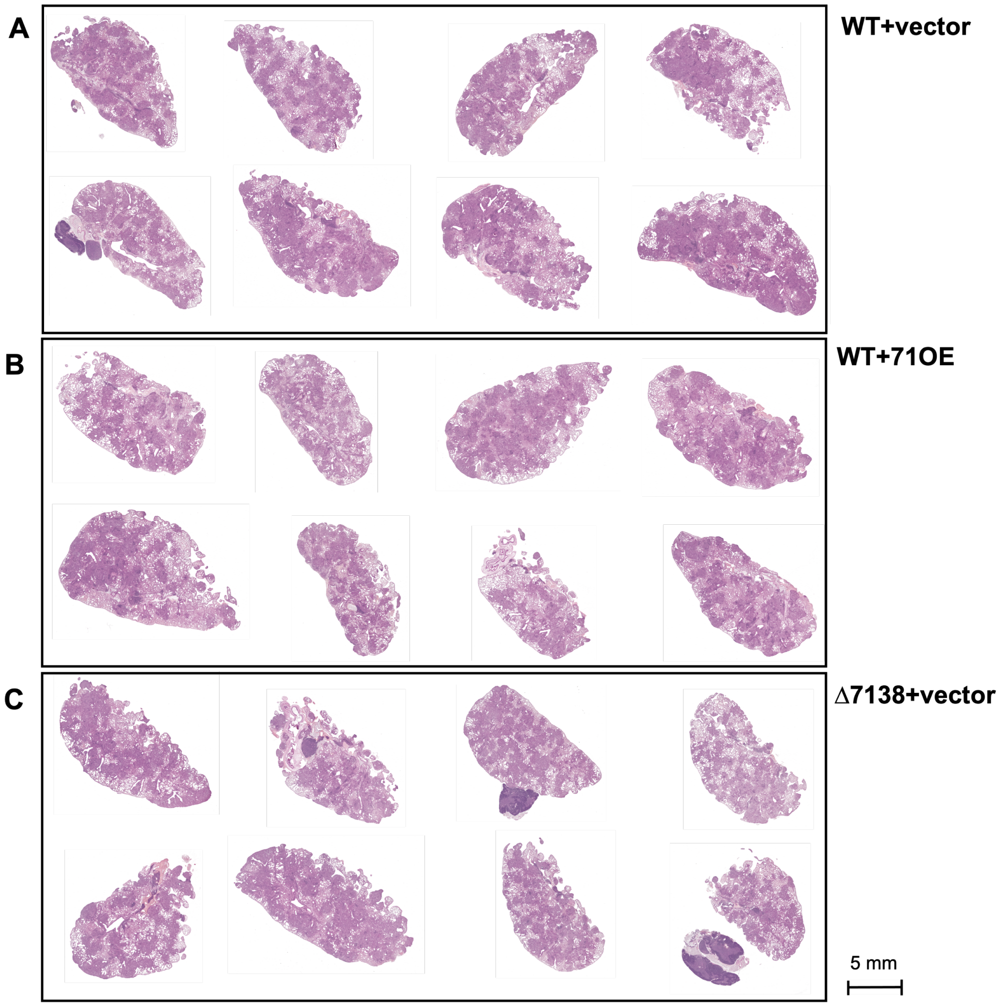


**Figure S2: Complete set of Week 8 lung histology images for WT+vector, WT+71OE, and Δ7138+vector mouse groups.**
(A–C) H&E histology images of lungs taken from female BALB/c mice infected with WT+vector (A), WT+71OE (B), or Δ7138+vector (C) strains at the Week 8 timepoint. These images were used to quantify the percent area of each lung occupied by inflammatory lesions. Non-pulmonary tissue present in the images (chiefly, upper airway and mediastinal lymph nodes) were excluded from this analysis.

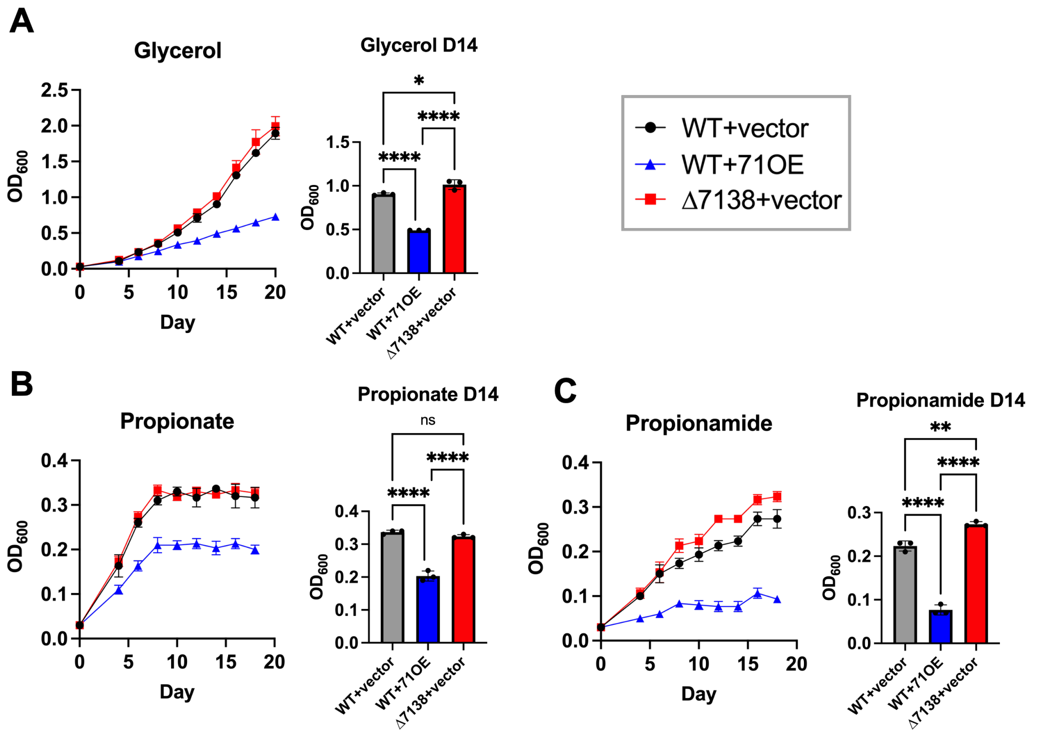


**Figure S3: H37Rv WT+71OE has a growth defect exacerbated by particular carbon sources.**

(A–C) Growth curves of *M.tb* strains in minimal 7H9 broth supplemented with 0.05% tyloxapol and each of (A) 0.2% glycerol, (B) 0.1% propionate, or (C) 5 mM propionamide. Graphs of optical densities at 600 nm ($OD_{600}$) for the Day 14 timepoints are provided for each medium. (mean±SD; n=3, ns: non-significant; *: $p<0.05$; **: $p<0.01$; ****: $p<0.0001$ by one-way ANOVA.)

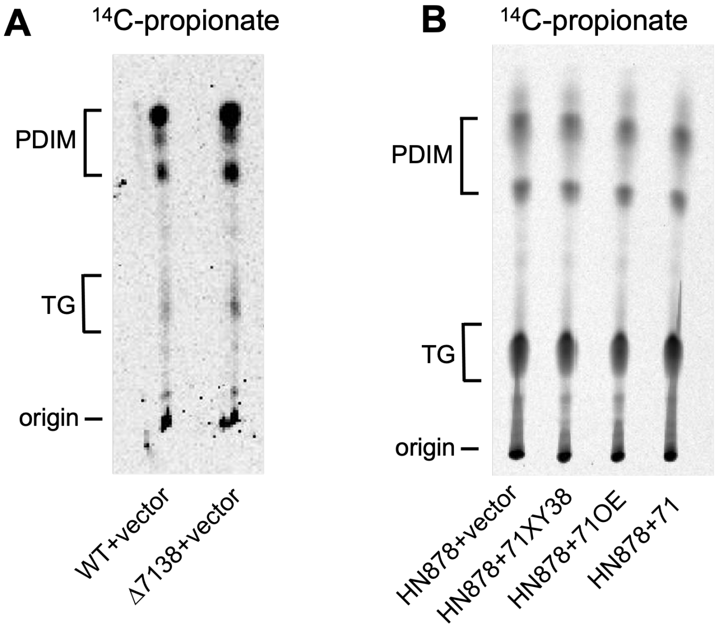


**Figure S4: M.tb H37Rv Δ7138 overproduces triglycerides.**
(A–B) TLC plates with apolar lipid fractions extracted from H37Rv WT+vector and Δ7138+vector strains (A) or HN878+vector, HN878+71XY38, HN878+71OE, and HN878+71 strains (B) labeled with $^{14}C$-propionate and visualized by phosphorimaging. Positions of phthiocerol dimycocerosate (PDIM) and triglyceride (TG) lipids, as well as the origin, are indicated.

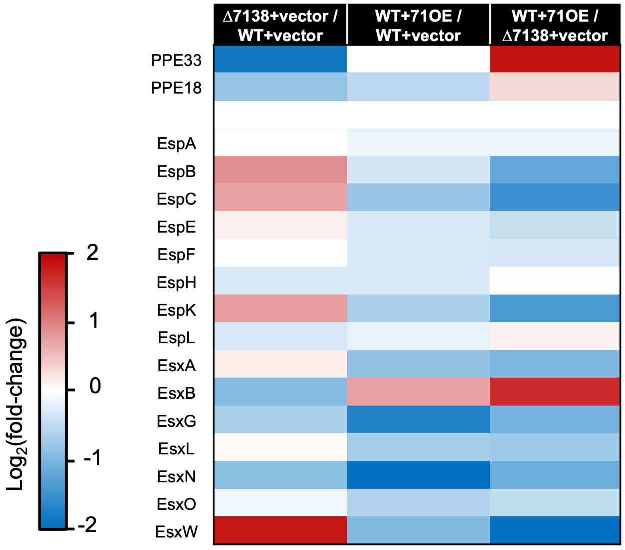


**Figure S5: Type VII secretion system substrates detected in culture filtrate.** Heat map of $\log_2$ fold-changes between each pair of strains for PE/PPE-, Esp-, and Esx- family proteins detected in CF fractions by mass spectrometry.

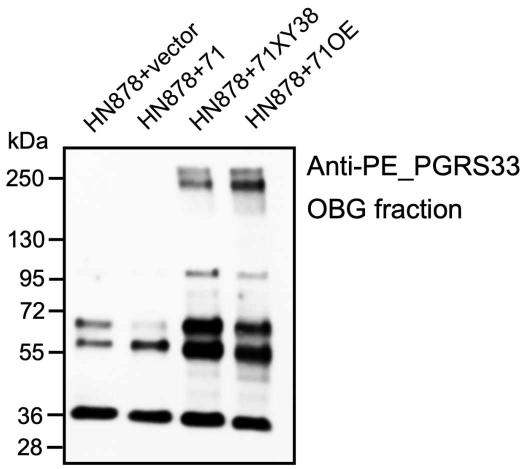


**Figure S6: Restoring the *PPE71* locus into HN878 strains boosts PE_PGRS protein secretion.**
HN878+vector, HN878+71, HN878+71XY38, and HN878+71OE cultures were incubated overnight in complete 7H9 broth without detergent. The octyl glucoside (OBG) fraction was subjected to Western blotting with anti-PE_PGRS33 antibody. Additional bands seen in the HN878+71XY38 and HN878+71OE strains suggest the presence of a broader range of PE_PGRS proteins. (kDa: kilodaltons.)

### Supplementary Tables

**Table S1. RNAseq output for transcripts from WT+vector, WT+71OE, and Δ7138+vector strains.**
Raw counts and normalized counts (scaled by total transcripts per sample) are provided. Fold-changes and adjusted p-values (using a Benjamini-Hochberg multiple test correction) were computed for each comparison.

**Table S2. Mass spectrometry output for the culture filtrate fraction of WT+vector, WT+71OE, and Δ7138+vector strains.**
Raw input and quantified proteins, normalized to total peptides per sample by summed signal/noise (S/N) ratio, are provided. Fold-changes and p-values were computed for each comparison.

**Table S3. Mass spectrometry output for the octyl glucoside fraction of WT+vector, WT+71OE, and Δ7138+vector strains.**
Raw input and quantified proteins, normalized to total peptides per sample by summed signal/noise (S/N) ratio, are provided. Fold-changes and p-values were computed for each comparison.

**Table S4. Plasmids used in this study.**

| **Plasmid** | **Description** | **Reference** |
|---|---|---|
| pMH94 | L5 *int*, *M.smegmatis attP*, *ampR* (*E.coli*), *kanR* (*M.tb*) | (71) |
| pMH94-PPE71 | pMH94 + [422nt upstream of *PPE71* to 111nt downstream of *PPE71*] | This work |
| pMH94-71XY | pMH94 + [414nt upstream of *PPE71* to 177nt downstream of *esxY*] | This work |
| pMH94-71XY38 | pMH94 + [414nt upstream of *PPE71* to 244nt downstream of *PPE38*] | This work |
| pSD5 | *OriM*, *M.leprae hsp65* promoter, *ampR* (*E.coli*), *kanR* (*M.tb*) | (72) |
| pSD5-71OE | pSD5 + [60nt upstream of *PPE71* to 28nt downstream of *PPE38*] | This work |
| pET-28a(+) | T7 promoter, lac operator, 6xHis-insert-6xHis, *kanR* (*E.coli*) | Novagen |
| pET28a-PPE71 | pET-28a(+) + *PPE71* [in frame with both N-terminal and C-terminal 6xHis] | This work |

**Table S5. Oligonucleotides used in this study.**

| Oligonucleotide | Sequence (5’–3’) |
|---|---|
| PPE71_24f | GAATTCGAGCTCGGTACCCGGGGATCCTCTAGATTCCGTTCGGTAGTGCGAT |
| PPE71_24r | GCAGAGATGGTGCCCTTGGTGGTCGACTCTAGTCATAAACCGAGTAGCCACCA |
| PPE71locus_59_F | GACCTCGGTACCAGGTAGTGCGATGTAGTTGGTCT |
| EsxY_59_R | GTCGACTCTAGACTCAACGGCTCAGACACAAAC |
| PPE38term_59_R | GTCGACTCTAGATCCTTCAGATCGCGATGGTTG |
| PPE71_pSD5_f | GCGATATCCGGAGGAATCACTTCCATGTTTGTGTCTGGAGAGTGGTAGG |
| PPE71_pSD5_r | CCATTGAAGACCGGGCCAGAACGCGGCAAAGACCCCGACCAATC |
| dPPE38_57_f | GTCACTAGACAGTTCGAGGTCA |
| dPPE38_57_r | GGTAGTGCGATGTAGTTGGTCT |
| PPE71_34F | ACCGAAAACCTTTACTTCCAGGGCCATATGTGGAGAGTGGTAGGCGGA |
| PPE71_34R | TGCTCGAGTGCGGCCGCAAGCTTGTCGACCCAATCACCTCCGCCGTATCC |
| pMH94_Fseq | GAAAATACCGCATCAGG |
| pMH94_Rseq | GAATAGACCGGGACAAGG |
| PPE71-EsxX_Fseq | GTCTTTGCGTTGATGACAT |
| PPE71-EsxX_Rseq | ATGTCATCAACGCAAAGAC |
| EsxY-PPE38_Fseq | CCTTCGGCATGTCAACA |
| EsxY-PPE38_Rseq | TGTTGACATGCCGAAGG |
| pSD5_insert_FWD | AGCGTAAGTAATGGGGGTTGTCG |
| pSD5_insert_REV | ATATATTCGGTCGCTGAGGCTTG |
| dPPE38_inner_f | CAGCGTCCGTACCATG |
| dPPE38_inner_r | TTCGCGCAGTCTTTACG |
| T7_FWDseq | TAATACGACTCACTATAGGG |
| T7_REVseq | GCTAGTTATTGCTCAGCGG |
| 16S_q_F | GCCGTAAACGGTGGGTACTA |
| 16S_q_R | TGCATGTCAAACCCAGGTAA |
| PPE71_q_T1F | ATCACCGCATCAAACGAGGA |
| PPE71_q_T1R | TGGATTTTTCGTGGTTGCCG |
| EccC5_qF | AGGCTTGGTTCTTCGCTCTC |
| EccC5_qR | AGATGTCTCGGGCACCAATG |
| pfkB_q_1F | TTTCCCAAGCGAACGGACTT |
| pfkB_q_1R | GCGAGCCATCGATTTTCGTC |
| devS_q_9F | CCGAGTGATACCCTGCGATG |
| devS_q_9R | CCGCACGAACGTAATCCTGA |
| devR_q_5F | CGCCAACTCCATTCCCTTGA |
| devR_q_5R | GCTGTCTGATCCTCACGTCC |
| PPE51_q_2F | CGCAGGATGTCGAGTCCTTT |
| PPE51_q_2R | AACTCGCCGATCCCAAAGTC |
| radA_q_1F | GTCGACGTTGTGCTGCATTT |
| radA_q_1R | GTGCAGGAGGAAACATCCGA |
| hspX_q_1F | TGGACCGGATCTGAATGTGC |
| hspX_q_1R | CCACCTACGACAAGGGCATT |
| narX_q_4F | ATAGTCGGTCTCCTGCGTCT |
| narX_q_4R | TCCACACACGGGGTGAATTG |
| kasA_q_6F | ATGCTCATCGAGACGGAGGA |
| kasA_q_6R | ATGAAAGGCGTCCGAGGTG |
| fas_q_1F | GCCCCATCATCTGGAAGACC |
| fas_q_1R | CATCTCCAAGCACGACACCT |
| tgs1_q_3F | GCAGGTTGGGCAGCATTAAC |
| tgs1_q_3R | TCAATGATGTTGCGCTTGCC |
